# Proinflammatory S100A9 Regulate Differentiation and Aggregation of Neural Stem Cells

**DOI:** 10.1101/2020.06.06.137745

**Authors:** Yin Tian, Rui Cao, Bingchen Che, Yong Tang, Lin Jiang, Bai Qiao, Yonggang Liu, Ludmilla A Morozovaroche, Ce Zhang

**Author notes:** These authors contribute equally to this work (Yin Tian, Rui Cao).

## Abstract

Inflammation is the primary pathological feature of neurodegenerative diseases such as Alzheimer’s (AD) and Parkinson’s disease. Proinflammatory molecules (e.g. S100A9) play important roles during progression of the diseases by regulating behavior and fate of multiple cell types in the nervous system (*1*). Our earlier studies reveal that S100A9 is toxic to neurons, and its interaction with A*β* peptides leads to the formation of large non-toxic amyloidogenic aggregates, suggesting a protective role of A*β* amyloids (*2*). We herein, demonstrate that S100A9 interacts with neural stem cells (NSCs) and causes NSC differentiation. In the brain of transgenic AD mouse models, we found large quantities of proinflammatory S100A9, which colocalizes with the differentiated NSCs. NSC sphere formation, which is a representative character of NSC stemness, is also substantially inhibited by S100A9. These results suggest that S100A9 is a representative marker for the inflammatory conditions in AD, and it promotes NSC differentiation. Intriguingly, in contrast to the death of both stem and differentiated NSCs caused by high S100A9 doses, S100A9 at a moderate concentration is toxic only to the early differentiated NSCs (i.e. progenitor cells and immature neurons), but not the stem cells. We therefore postulate that at the early stage of AD, expression of S100A9 leads to NSC differentiation, which remedies the neuron damages. The application drugs, which help maintain NSC stemness (e.g. PDGF), may help overcome the acute inflammatory conditions and improve the efficacy of NSC transplantation therapy.

## 1 Introduction

Neural stem cells are the self-renewing, multipotent cells that are capable of unlimited proliferation and differentiating into phenotypes such as neurons, astrocytes and oligodendrocytes (*3–5*). When neuron loss occurs during acute and chronic nervous system diseases such as traumatic brain injury and neurodegenerative diseases, neurogenesis is expected to be triggered to mediate the damage (*6–8*). Numerous physiological and pathological environmental factors including stress, aging and inflammation can affect NSC behavior and contribute to the fact that neurogenesis happens mostly in the young adult brain but not in the aged brain (*9–11*). A*β* amyloidogenic species, which are the hallmark of AD, are found to have distinctive effect on NSCs (*12–15*). For example, oligomeric A*β*_42_ was found to promote NSC differentiation, but not monomeric and fibrillary A*β*_42_ (*16–21*). It is proposed that since A*β*_42_ fibrils are the prevalent aggregation state in the end-stage of AD, NSC differentiation when being in contact with A*β*_42_ oligomers may be a defense mechanism of the nervous system (*22*).

Another major pathological symptom of AD, which is closely related to both neuron death and A*β* amyloid formation, is the overexpression of proinflammatory S100A9. In nervous system, S100A9 is mainly expressed by activated microglial cells and endothelial cells, and associated with several neurodegenerative diseases (*23–28*). As one of the calcium-binding proteins, S100A9 takes on diverse roles during life machinery, including cell proliferation, migration, invasion, stress and differentiation (*29–38*). In AD, S100A9 and its family proteins are found to co-localize with senile plaques, and serve as key mediator of amyloid cytotoxicity (*39*). Our earlier studies reveal that under pathological conditions, S100A9 actively interact with Aβ_42_, and the resulting amyloidogenic aggregates may act as a sink for the toxic species. Through numerical simulation, we further demonstrate the savior effect of S100A9 is related to its characteristic structural features. However, the effect of S100A9 on NSCs remain unclear.

In this work, we explore the effect of S100A9 on NSCs by monitoring the expression level of Hes5, Dcx, nestin and cellular behavior including differentiation and proliferation. We demonstrate that S100A9 promotes NSC differentiation, and subsequently cause cytotoxicity on early neurons originated from NSCs. Our studies reveal a new mechanism of AD development, in which the stem cell pool is firstly exhausted by inducing NSC differentiation, and later killed by S100A9. As NSC implantation is proven to be a promising strategy in regeneration medicine, the identification of pathological conditions, which may affect the residency of NSCs, is of great importance for therapeutic applications.

## 2 Results

### Overexpression of S100A9 leads to NSC differentiation in APP/PS1 mice

In the hippocampus of APP/PS1 mice, we observe that S100A9 expression level is significantly higher than that in control group (C57bl/6 mouse) (Fig. 1a-d). The expression level of S100A9 of 6-month and 8-month old APP/PS1 mice is ~40% and ~50% higher than the control samples, respectively (Fig. 1e). There is no significant difference before 4-month-old APP/PS1 mice as compared to the control group. With aging, the S100A9 level in the hippocampus region of APP/PS1 mice nearly doubles from 4 to 8 months (Fig. 5f), and remain unchanged in control mice. Notably, S100A9 molecules largely co-localize with Dcx-positive cells in the CA3 region of hippocampus (Fig. S1), suggesting effect of S100A9 on NSC differentiation.

**Figure 1:**
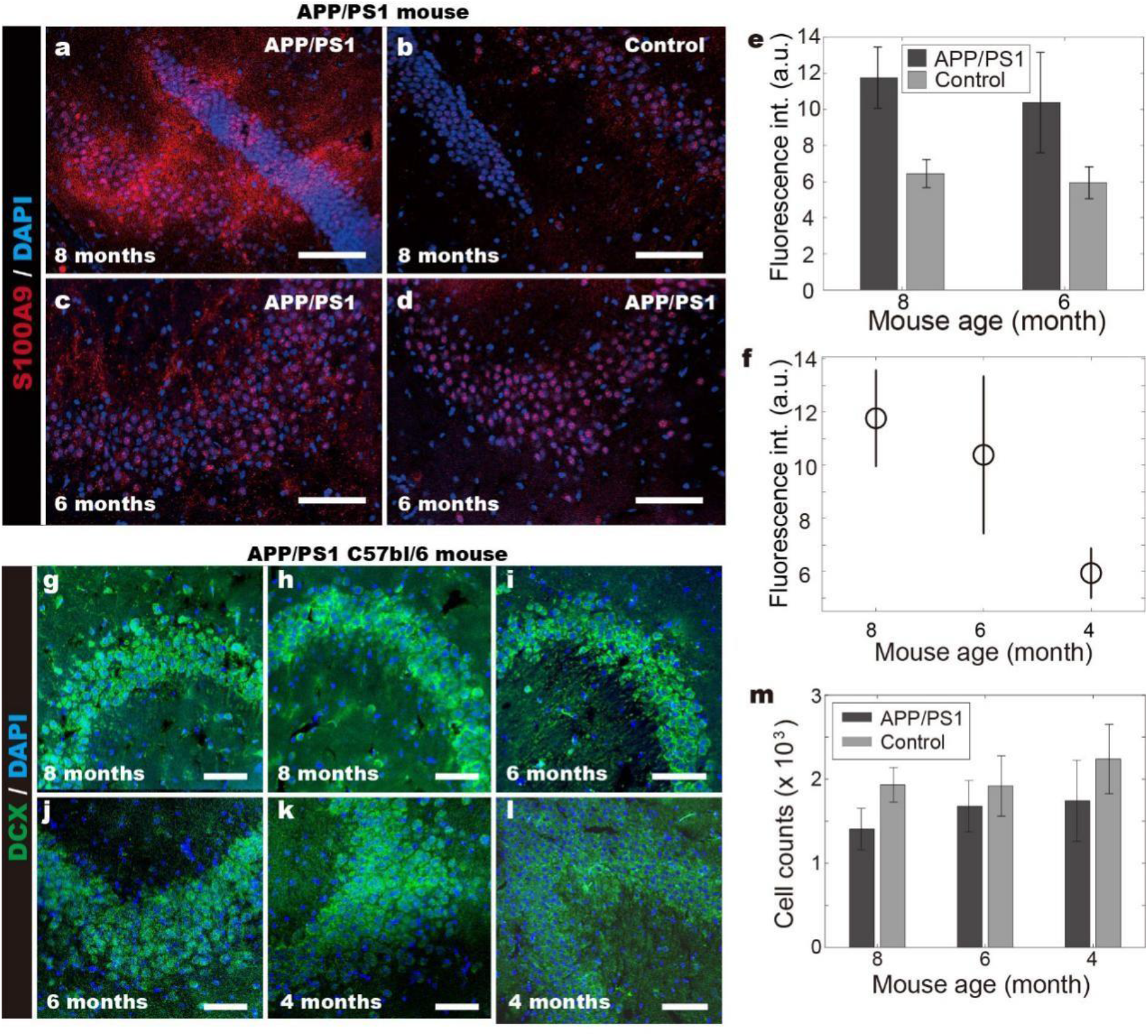
Increased expression of S100A9 in hippocampus of APP/PS1 mice. **(a,b)** 8-month-old a. APP/PS1, and b. control C57bl/6 mouse hippocampus. **(c,d)** 6-month-old c. APP/PS1, and d. control C57bl/6 mouse hippocampus. **(e,f)** Expressionlevels of S100A9 in hippocampus of APP/PS1 and control mice at different days. **g.** 8-month-old APP/PS1 C57bl/6 mouse hippocampus. **h.** 8-month-old C57bl/6 mouse hippocampus. **i.** 6-month-old APP/PS1 C57bl/6 mouse hippocampus. **j.** 6-month-old C57bl/6 mouse hippocampus. **k.** 4-month-old APP/PS1 C57bl/6 mouse hippocampus. **l.** 4-month-old C57bl/6 mouse hippocampus. **m.** Statistical results showing S100A9 induced death of neurons and early differentiated cells. Scale bars denote 85 *μ*m in **a-d**, and 120 *μ*m in **g-l**.

By counting the number of Dcx-positive cells, we observe that the number of neurons and early differentiated NSCs inversely correlate with S100A9 concentration (Fig. 1f and 1m). Besides the decreased cell counts, the expression level of Dcx increases as well, suggesting differentiation (Fig. 1g-l). It is demonstrated that despite of the large variances from 4 months to 6 months, which may be caused by the compensation of adult neurogenesis in hippocampus (*40*), the number of Dcx-positive cells remains nearly unchanged, from 6 to 8 months in the control samples (Fig. 1m). In contrast, the quantities of Dcx-positive cells in APP-PS1 mice keep decreasing. These results suggest that overexpression of S100A9 leads to the death of neurons and early differentiated NSCs, which are Dcx-positive (*41*).

### S100A9 suppresses NSC aggregation and promotes differentiation

Hes5 is an essential factor of Notch signaling pathways, which regulate the maintenance of NSC stemness and repress neural differentiation (*42, 43*). We demonstrate that NSC sphere formation is initiated by physical contact of cell-cell and cell-axon (*44*), upon which the Hes5 fluorescent signal increases. Meanwhile, the development of axons, which direct NSC migration, is not limited by the laminin-coating (Video S1). During both adherent and suspension culture, NSC spheres actively move and recruit resident cells. It is worth noting that the organization of spheres is highly dynamic. Cells constantly detach themselves, immediately after which the Hes5 level drops to their original value before cell-cell contacts (Fig. 2c and Video S2). These results suggest that Hes5 is not only involved in Notch signaling upon physical contact with other cells or axons, it may help regulate cell aggregation, where single NSC cells are grouped based on the Hes5 level. The hypothesis is consistent with what we observed during continuous culture of NSC spheres with both Hes5 and Dcx fluorescent markers. It is demonstrated that the cells with high Dcx and low Hes5 level, which reflect differentiation (*41–43*), locate themselves on the outer layer of the sphere. While, cells with high Hes5 and low Dcx level merge into an entity, in which no distinguishable boundaries can be observed (Video S3).

**Figure 2:**
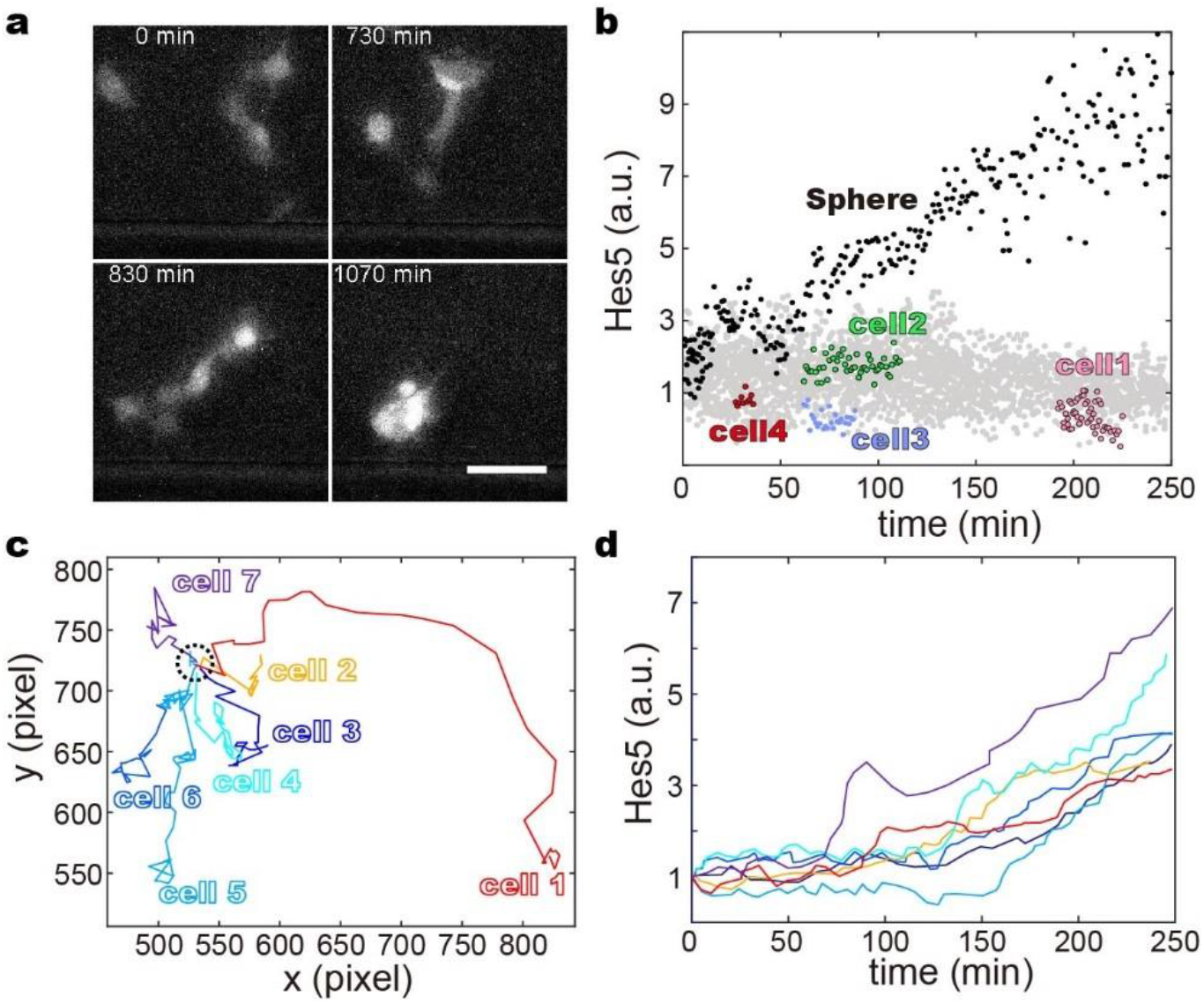
NSC sphere formation associates with Hes5 expression level. **a.** Variations in NSCs Hes5 level during sphere formation. **b.** NSC cells included in spheres (dark) and the stand-alone ones (grey) show different Hes5 level. We observe that NSCs with low Hes5 level (cell 1-4) dissociate themselves from sphere. **c.** Trajectories of NSCs cells before aggregation. **d.** Hes5 level individual cells before and after NSC sphere formation.

The application of S100A9 inhibits NSC sphere formation (Fig. 3b-f), and down-regulates nestin expression (Fig. 3g), which is a representative marker for NSCs stemness (*45–47*). In the meantime, the Hes5 level remain unchanged during 2 days’ culture, suggesting inactivation of Notch signaling pathway due to the lack of cell-cell and cell-axon contact (Fig. 2 and Fig. 3g). Intriguingly, S100A9 at the dose of 10 *μ*g/ml, which induces ~ 70 % death rate of Dcx-high cells, has no substantial effect on the viability of NSCs with high Hes5 level (Fig. 3h). Significant NSC cell death is observed when S100A9 concentration reaches 40 *μ*g/ml. These results suggest that S100A9 does not directly cause NSC cell death. Instead, it inhibits Notch signaling by preventing the physical contacts among NSC cells, which plays important roles in the maintenance of stem cells (*48–52*). Subsequently, NSCs differentiate into progenitor cells or neurons with prolonged incubation (days), which is considerably more vulnerable to S100A9 stimulation.

**Figure 3:**
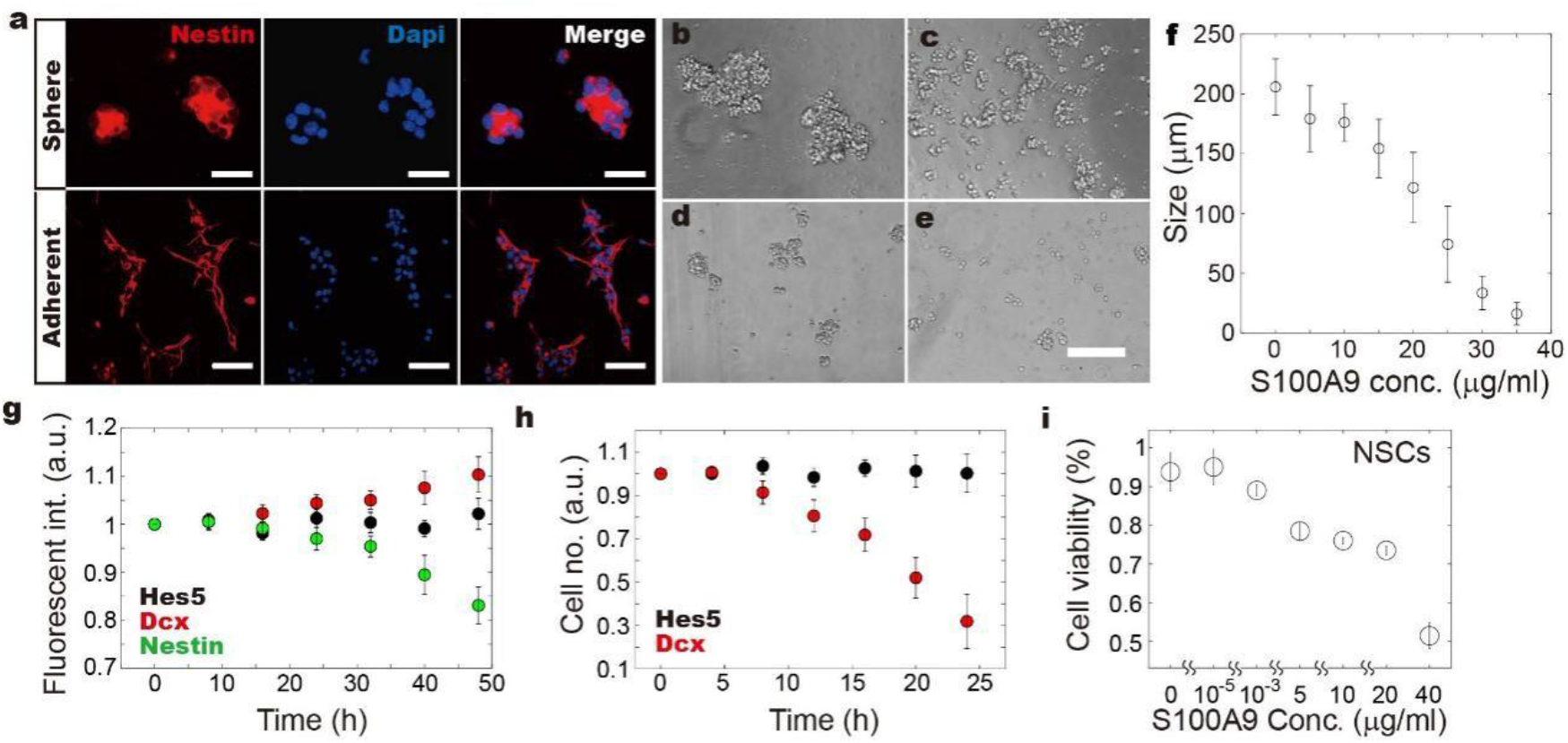
S100A9 affect NSC sphere formation and the expression level of Dcx, Hes5 and Nestin. **a.** NSC sphere and adherent cells showing variations in nestin expression level. **b-e.** Bright-field images of NSCs maintained as suspension cells, and incubated with different doses of S100A9 for 24 h, **b.** 0 *μ*g/ml; **c.** 10 *μ*g/ml; **d.** 20 *μ*g/ml; **e.** 40 *μ*g/ml. We observe that S100A9 inhibits NSC aggregation, and thus NSC sphere formation, which is known to be one of the characteristics representing stemness (*53*). **f.** The size of NSC spheres decreases with increasing S100A9 concentration. **g.** Fluorescence level of Hes5, Dcx and Nestin upon stimulation by 10 *μ*g/ml S100A9. **h.** Counts of cells with high Hes5 (dark) and high Dcx (red) level during incubation with 10 *μ*g/ml S100A9. **i.** Variations in cell viability depends on S100A9 concentration.

### S100A9-induced NSC differentiation is mediated by PDGF

Analysis of RNA sequencing data reveals up- and down-regulated genes upon S100A9 stimulation (Fig. 4a). As S100A9 inhibits NSC aggregation, we firstly study the variations in genes related to cell migration and axon development. It is demonstrated that the genes related to cytoskeleton reorganization (i.e. Map2, Atat1, Tubb3, Capza1 and Capzb) is not significantly affected (*54–57*). In contrast, genes related to axon guidance are mostly down-regulated, suggesting a cause and consequence relationship between S100A9 stimulation and axon development. We suggest that the variations in genes, which are associated with or activated upon the physical contacts among cells (e.g. cell attachment, cell adherent molecules (CAMs) and Notch signaling), are caused by prohibited NSC aggregation.

**Figure 4:**
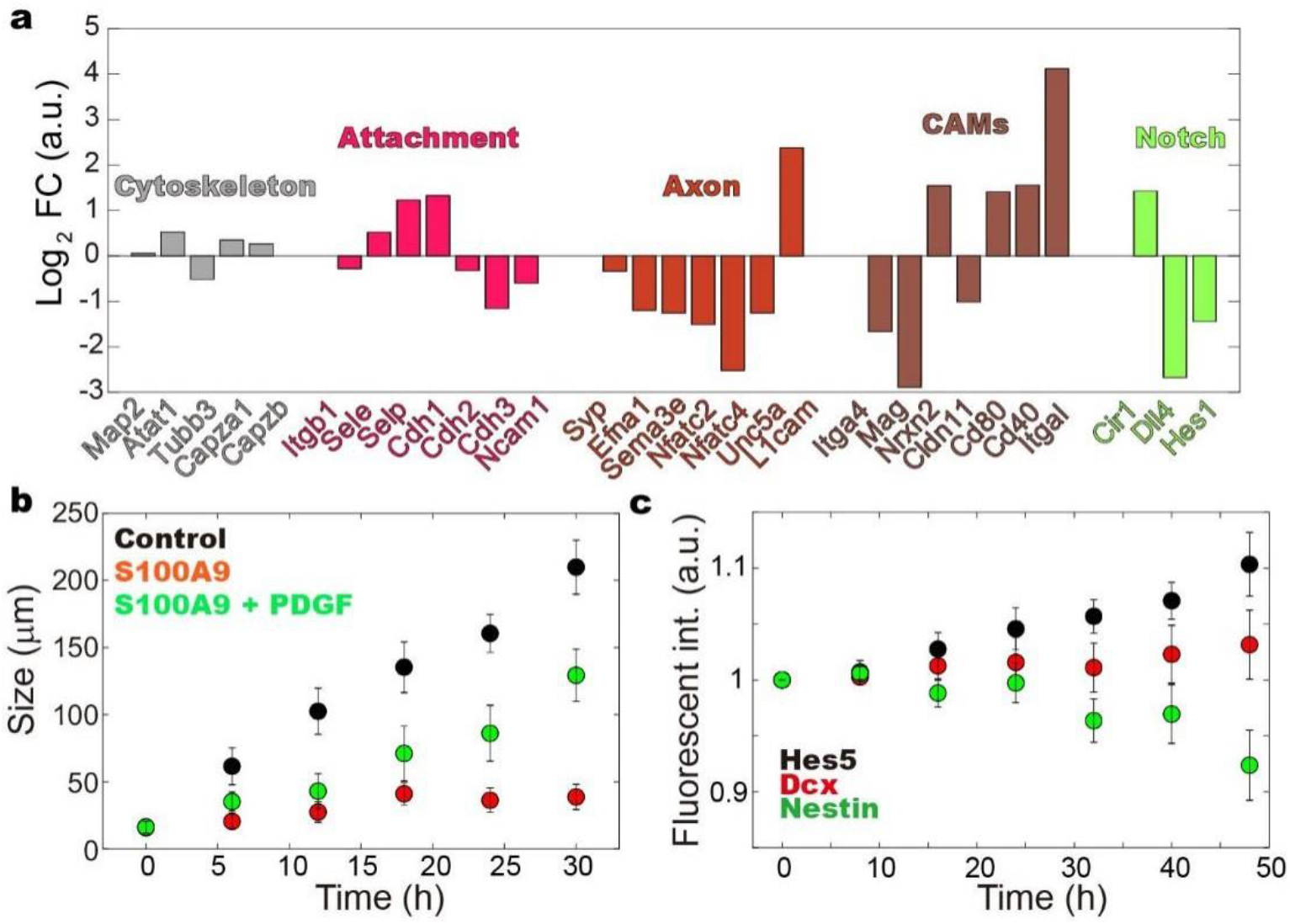
Variation in gene expression level upon stimulation by S100A9. **a.** Analysis of RNA sequencing data reveals up- and down-regulated genes upon 10 *μ*g/ml S100A9 stimulation (see Supplementary Fig. 2 and Table 1), which are associated with NSC migration, and physical contact. **b.** The addition of 1 *μ*g/ml PDGF-AA (green) mediate the effects caused by 20 *μ*g/ml S100A9 (red). **c.** The variations in Hes5, Dcx and Nestin level is caused by NSCs incubation with 1 *μ*g/ml PDGF-AA and 20 *μ*g/ml S100A9.

To mediate the effect of S100A9 on NSCs, platelet-derived growth factor (PDGF-AA), which is known to promotes proliferation, survival and migration in diverse cell types, is introduced to the cellular microenvironments (*58–60*). It is demonstrated that the addition of PDGF simultaneously with S100A9 restores the capacities of NSC sphere formation (Fig. 4b). In the meantime, PDGF brings ~10% increase in the nestin level as compared to the samples co-incubated with only S100A9 for 2 days (Fig. 4c). The unchanged Dcx fluorescence signal and increase in Hes5 level by ~ 10% suggest well-maintained NSC stemness, and possibly the activation of Notch signaling (*61*).

## 3 Discussion

We herein demonstrate that NSC aggregation is initiated by establishing axon-cell contacts, which then direct cell and sphere migration. Addition of S100A9 down-regulates expression of the genes associated with axon development, and inhibits NSC sphere formation (Fig. 4a). Notch signaling, which plays important roles in the maintenance of NSC stemness, is down-regulated. As a result, NSC differentiate into progenitor cells and immature neurons, which show elevated Dcx and unchanged Hes5 level (Fig. 3g). Notably, S100A9 induces significant cell death mostly among the differentiated cells (high-Dcx), but not the stem cells (high-hes5), indicating a regulatory role of the inflammatory signals (Fig. 3h) (*62*). It is possible that at the early stage of neurodegenerative diseases, the moderate S100A9 concentration induces NSC differentiation to compensate the loss of neurons. The further increases in S100A9 causes the death of neurons, and eventually NSC cells (Fig. 3i). As reported in our earlier studies, the cytotoxicity of S100A9 can be mediated by A*β* peptides (*2*), resulting in the formation of large amyloidogenic aggregates.

We further demonstrate that NSC differentiation and cell death caused by S100A9 can be mediated by PDGF-AA, which is reported to affect NSC proliferation, migration, recruitment, and fate by working closely with Notch signaling (Fig. 4b-c) (*63*). We therefore suggest that Notch signaling is heavily involved in the regulatory process of both S100A9 and PDGF. The competition between positive and negative pathways leads to mediated S100A9 effect on NSCs. As the inflammatory conditions are the major symptoms of neurogenerative diseases (*22, 23*), studies on the effect of pro-inflammatory molecules (e.g. S100A9) on stem cells have great therapeutic potential as well. For example, stem cell transplantation as treatment for neurodegenerative diseases, can be affected by the level of S100A9 in the brain (*24*), in which PDGF-AA can be used as a drug ensuring NSC stemness.

## Materials and methods

### Animals and chemical materials

APP/PS1 double transgenic mice were provided by the Animal Model Institute of Nanjing University and C57bl/6 mice were provided by the Experimental Animal Center at Chongqing Medical University, P. R. China. ALL animals used in this study were maintained in strict accordance with the approval granted by Chongqing Municipal Committee of Science and Technology (SYXK/YU 2018-0003). The employed primary antibodies include: Nestin (rabbit polyclonal,1:200, Abcam), S100A9 (rabbit polyclonal, 1:100, Abcam), DCX(goat polyclonal, 1:200, Abcam), Hes5 (rabbit polyclonal, 1:200, Abcam); secondary antibodies include: DyLight 549, Donkey Anti-Rabbit IgG and DyLight 488, Donkey Anti-Goat IgG reagent kits (AmyJet Scientific). DAPI (Genview) was used to stain the nuclears. All chemicals were purchased from Sigma except those mentioned in this article.

### Isolation and culture of Neural stem cells

Neural stem cells (NSCs) were isolated and cultured from day 16 SD embyro-murine forebrain and Hes5-GFP/DCX-Red double transgenic mice. After rinsing in phosphate saline buffers (PBS), dissected cortical hemispheres were sliced into small pieces in icecold D-hank’s (Solarbio), which then were mechanically and enzymatically dissociated by the Papain Dissociation System as indicative materials (Worthington Biochemical Corp). The primary dissociated cells were maintained in neurobasal medium (Cyagen Biosciences) containing 2% (v/v) NS21 supplement, 0.5 mM L-glutamine, 100 units/ml penicillin and 100 mg/ml streptomycin under culture conditions, i.e. 95% humidity, 37°C and 5% (v/v) CO_2_. Neurospheres were formed within 1 week after the initial planting with semi-volume neurobasal medium exchange every 2-3 days. The neurospheres were passaged every 5-7 days by mechanically and enzymatically dissociation.

### Immunofluorescence staining of cells

For immunostaining, neurospheres and adherent cells were fixed in 4% polyparaformaldehyde (PFA) for 15 minutes at room temperature. The cells were then rinsed repeatedly (at least 3 times) with PBS for 5 mins. After pre-incubation in PBS containing 5% normal goat serum and 0.3% Triton X-100 for 15 min., samples were treated with the primary antibodies and incubate overnight at 4°C. Association of secondary antibodies was performed after thoroughly rinsing in PBS, which was followed by incubation in dark at 37 °C.

### Tissue processing

Being anesthetized via an intraperitoneal injection of 1% pentobarbital sodium (each mouse for 0.4 ml/100 g), a perfusion needle was planted into the exposed left ventricle, and the right atrial appendage was removed. After perfusion with saline solution, the mouse were then perfusion-fixed by 2% paraformaldehyde and 2.5% glutaraldehyde in 0.1 M PBS (pH 7.4). The olfactory bulb and cerebellum were removed from the perfused brain, remainder of which was dissected into two hemispheres. Gradient dehydration of the cerebral hemisphere in 10%, 20%, 30% sucrose solution for 24 h > 24h > 36h respectively. Randomly selected hemispheres were sliced into serial sagittal plane sections with a thickness of 10 μm using a cryo-ultramicrotome (Leica, Germany), which were then stained by immunofluorescence.

### Immuno-staining of hippocampus slices

For immune-staining, the frozen sections were rinsed twice with 0.1 M PBS solution for 30 min, and then with 0.3% Triton X-100. The slices were placed in citrate solution, and maintained in boiling water bath for 30 min to repair antigen. For the binding of primary antibodies, the slices were firstly incubate in 0.3% Triton X-100 containing 0.1% cold water fish gelatin, 1% bovine serum albumin and 5% normal goat serum for 2 h at 37°C, and for then 12 h at 4°C with primary antibodies against S100A9 and DCX. After incubation at 37°C for 1 h, the sections were maintained in solution for secondary antibodies for 2 h. Fluorescent markers were added to the solution, and incubated for 15 min at room temperature. Finally, the sections were mounted on gelatin-coated slides with antifade solution to reduce fluorescence quenching.

### Image acquisition and data analysis

The real-time fluorescent images of cells, tissues and biopsy samples were obtained using Ti2E inverted microscope and A1 confocal microscope (Nikon, Japan). Using customized Matlab program, image of the whole hippocampus is divided into different equinoxes at single pixel accuracy. Expression level of S100A9, nestin, Hes5 and Dcx are assessed by averaging the fluorescence intensity within a single cell.

## Acknowledgement

Y. L. acknowledge support from the Chongqing Basic and Frontier Research Program Project (cstc2015jcyjA10034).

