## Supplementary Information for "Proinflammatory S100A9 Regulate Differentiation and Aggregation of Neural Stem Cells"

### Movie descriptions

**Supplementary Video 1:** NSCs sphere formation during suspension and adherent culture. Notably, the PDMS surface for adherent culture is treated with laminin for merely 20 min, which is considerably shorter than the standard protocol.

**Supplementary Video 2:** The NSCs sphere formation process is highly dynamic, involving constant cell migrating in and out the entity.

**Supplementary Video 3:** NSCs cells seems to group themselves based on the Dcx and Hes5 level during sphere formation and continuous culture. The cell with high Dcx level distribute only on the outer layer of the sphere, and the ones with high Hes5 stay in the core region.

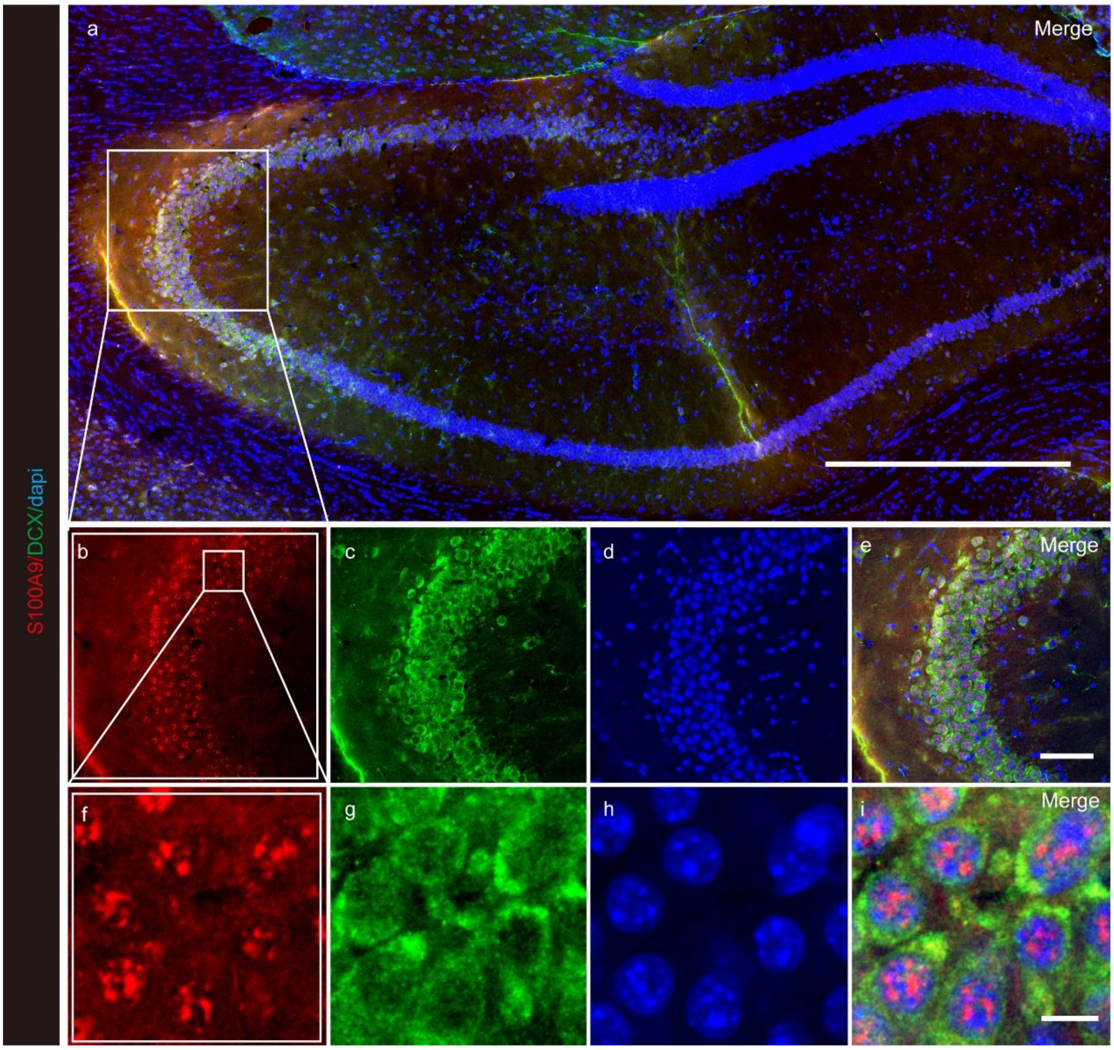


**Supplementary Figure 1** S100A9 and DCX$^{+}$ cells co-localization in the hippocampus. **a.** hippocampus large scan, Scale bars, 500 μm. **b-e.** Partial enlargement of hippocampal CA3 area, Scale bars, 65 μm. **f-I.** Show details, Scale bars, 10 μm.

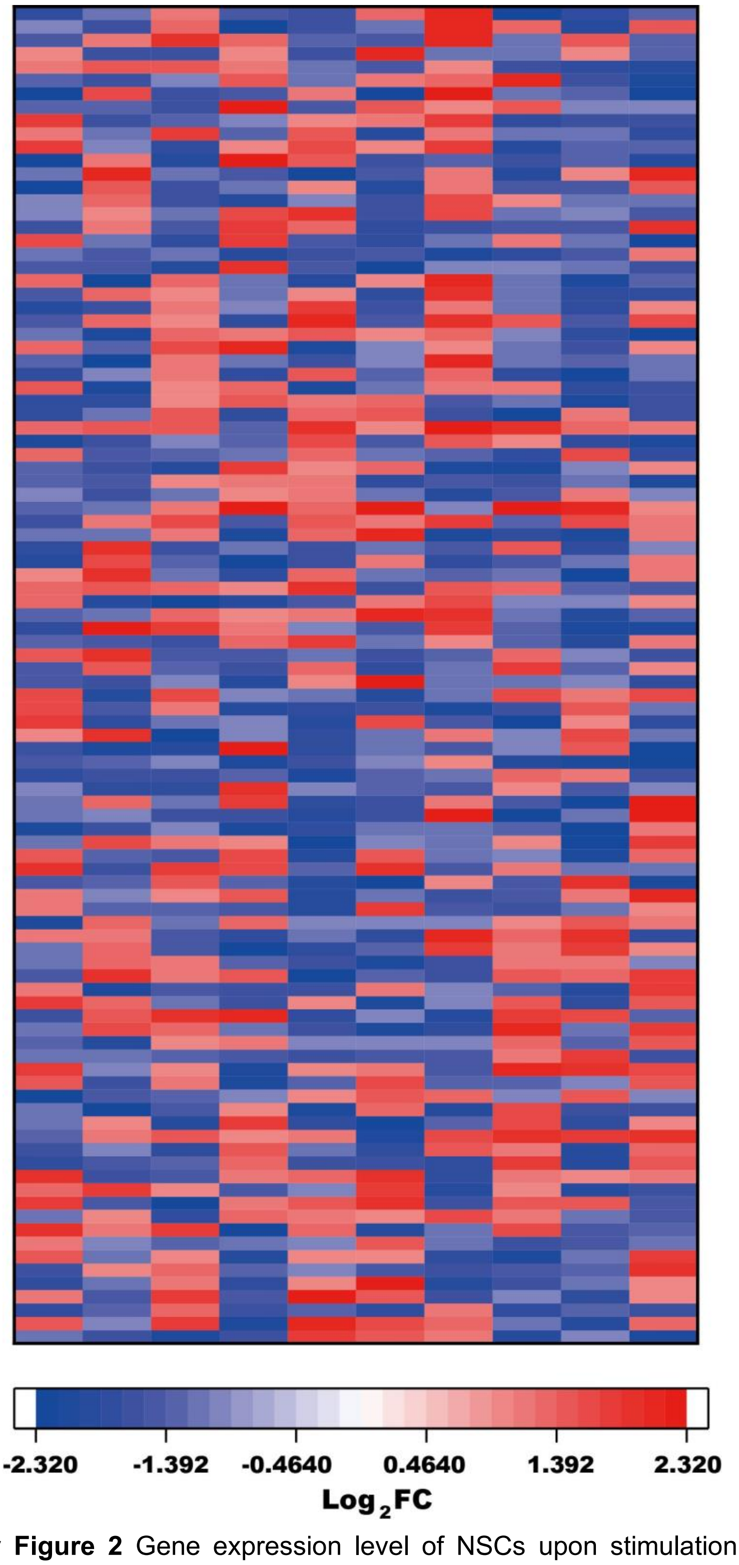


**Supplementary Figure 2** Gene expression level of NSCs upon stimulation by S100A9 is normalized to the values of control samples, which are maintained in the culture medium. It is demontrated that addtion of 10 µg/ml S100A9 causes substantial amount of up- and down-regulated genes. The list of gene names is shown in Supplementary Table 1.
